# A Forward Model at Purkinje Cell Synapses Facilitates Cerebellar Anticipatory Control

**DOI:** 10.1101/078410

**Authors:** Ivan Herreros-Alonso, Xerxes D. Arsiwalla, Paul F.M.J. Verschure

## Abstract

How does our motor system solve the problem of anticipatory control in spite of a wide spectrum of response dynamics from different musculo-skeletal systems, transport delays as well as response latencies throughout the central nervous system? To a great extent, our highly-skilled motor responses are a result of a reactive feedback system, originating in the brain-stem and spinal cord, combined with a feed-forward anticipatory system, that is adaptively fine-tuned by sensory experience and originates in the cerebellum. Based on that interaction we design the counterfactual predictive control (CFPC) architecture, an anticipatory adaptive motor control scheme, in which a feed-forward module, based on the cerebellum, steers an error feedback controller with *counterfactual* error signals. Those are signals that trigger reactions as actual errors would, but that do not code for any current of forthcoming errors. In order to determine the optimal learning strategy, we derive a novel learning rule for the feed-forward module that involves an eligibility trace and operates at the synaptic level. In particular, our eligibility trace provides a mechanism beyond co-incidence detection in that it convolves a history of prior synaptic inputs with error signals. In the context of cerebellar physiology, this solution implies that Purkinje cell synapses should generate eligibility traces using a forward model of the system being controlled. From an engineering perspective, CFPC provides a general-purpose anticipatory control architecture equipped with a learning rule that exploits the full dynamics of the closed-loop system.

## 1 Introduction

Learning and anticipation are central features of cerebellar computation and function [Bastian, 2006]: the cerebellum learns from experience and is able to anticipate events, thereby complementing a reactive feedback control by an anticipatory feed-forward one [Hofstoetter et al., 2002]. This interpretation is based on a series of anticipatory motor behaviors that originate in the cerebellum. For instance, anticipation is a crucial component of acquired behavior in eye-blink conditioning [Gormezano et al., 1983] where an initially neutral stimulus such as a tone or a light (the conditioning stimulus, CS) is followed, after a fixed delay, by a noxious one, such as an air puff to the eye (the unconditioned stimulus, US). As a result, during early trials, a protective unconditioned response (UR), a blink, occurs reflexively in a feedback manner following the US. After training though, a well-timed anticipatory blink (the conditioned response, CR) precedes the US. Thus, learning results in the (partial) transference from an initial feedback action to an anticipatory (or predictive) feed-forward one. Similar responses occur during anticipatory postural adjustments, which are postural changes that precede voluntary motor movements, such as raising an arm while standing. The goal of these anticipatory adjustments is to counteract the postural and equilibrium disturbances that voluntary movements introduce. These behaviors can be seen as the feedback reactions to events that after learning have been transferred to feed-forward actions anticipating the events.

Anticipatory feed-forward control can yield high performance gains over feedback control whenever the feedback loop exhibits transmission (or transport) delays [Jordan, 1993]. However, even if a plant has negligible transmission delays, it may still have sizable inertial latencies. For example, if we apply a force to a visco-elastic plant, its peak velocity will be achieved after a certain delay; i.e. the velocity itself will lag the force. An efficient way to counteract this lag will be to apply forces anticipating changes in the desired velocity. That is, anticipation can be beneficial even when one can act instantaneously on the plant. However, returning to the case of the cerebellum, the question is, what would be an optimal strategy to learn anticipatory actions?

To answer that we design the counterfactual predictive control (CFPC) scheme, a cerebellar-based adaptive-anticipatory control architecture that learns to anticipate performance errors from experience. With CFPC, we propose a generic scheme in which a feed-forward module enhances the performance of a reactive error feedback controller driving it with signals that facilitate anticipation, namely, with counterfactual errors. These are signals that drive the error feedback controller, enabling predictive control, that are learned based on experienced errors, but that do not reflect any actual or forthcoming error. Concretely, the CFPC scheme is motivated from neuro-anatomy and physiology of eye-blink conditioning. It includes a reactive controller, which is an output-error feedback controller that models brain stem or spinal cord reflexes actuating on eyelid or limb muscles, and a feed-forward adaptive component that models the cerebellum and learns to associate its inputs with the error signals derived from the signals driving the reactive controller.

In addition to eye-blink conditioning and postural adjustments, the interaction between reactive and cerebellar-dependent acquired anticipatory behavior has also been studied in paradigms such as visually-guided smooth pursuit eye movements [Lisberger, 1987] and predictive grasping [Nowak and Hermsdörfer, 2006]. All these paradigms can be abstracted as (predictable) reference tracking or disturbance rejection tasks that are trained in a trial-by-trial manner. That is, the same predictive stimuli and disturbance or reference signal are repeatedly experienced. In accordance to that, we operate our control scheme in trial-by-trial (batch) mode. With that, we derive a learning rule for anticipatory control that modifies the well-known least-mean-squares/Widrow-Hoff rule with an eligibility trace. More specifically, our model predicts that to facilitate learning, parallel fibers to Purkinje cell synapses implement a forward model that generates an eligibility trace. Finally, to stress that CFPC is not specific to eye-blink conditioning, we demonstrate its application with a smooth pursuit task.

## 2 Methods

### 2.1 Cerebellar Model

We follow the simplifying approach of modeling the cerebellum as a linear adaptive filter, while focusing on computations relevant to the Purkinje cells, which are the main output cells of the cerebellar cortex [Fujita, 1982, Dean et al., 2010]. Over the so-called mossy fiber input pathway, the cerebellum receives a wide range of inputs. Those inputs reach Purkinke cells via parallel fibers (Fig. 1), that cross dendritic trees of Purkinje cells in a ratio of up to 1.5 × 10^6^ parallel fiber synapses per cell. The signal carried by a particular fiber is indicated as *x*_*j*_, *j* ϵ [1, *G*], with *G* denoting the total number of inputs fibers. These inputs from the mossy/parallel fiber pathway carry contextual information (interoceptive or exteroceptive) that allows the Purkinje cell to generate a functional output. We refer to these inputs as *cortical bases*, indicating that they are localized at the cerebellar cortex and that they provide a repertoire of states and inputs that the cerebellum combines to generate its output *o*. As we will develop a discrete time analysis of the system, we use *n* to indicate time (or time-step). Hence, the output of the cerebellum at any time point *n* results from a weighted sum of those cortical bases. *w*_*j*_ indicates the weight associated with the fiber *j*, or synaptic efficacy. Thus, we have ***x***[*n*] = [*x*_1_[*n*],…,*x*_*G*_[*n*]]^*T*^ and ***w***[*n*] = [*w*_1_[*n*],…, *w*_*G*_[*n*]]^*T*^ (where the transpose, *T*, is meant to indicate that ***x***[*n*] are ***w***[*n*] column vectors) containing the set of inputs and synaptic weights at time *n*, respectively, which determine the output of the cerebellum according to

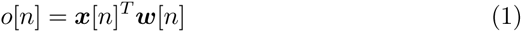

**Figure 1:**
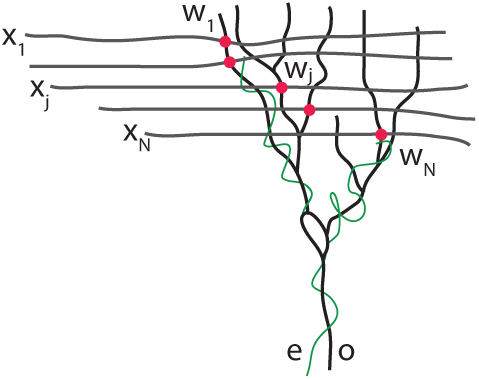
Anatomical scheme of a Cerebellar Purkinje cell. The *x*_*j*_ denote parallel fiber inputs to Purkinje synapses (in red) with weights *w*_*j*_. *o* denotes the output of the Purkinje cell. The error signal *e*, through the climbing fibers (in green), modulates synaptic weights.

The adaptive feed-forward control of the cerebellum stems from updating the weights according to a rule of the form

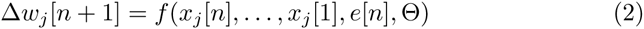

where Θ denotes global parameters of the learning rule. *w*_*j*_[*n*] represents the current efficacy of a “synapse” *j*, (*x*_*j*_[*n*],…, *x*_*j*_[1]), the history of its pre-synaptic inputs and *e*[*n*] an error signal that is the same for all synapses/weights, that corresponds to the difference between the desired, *r* and the actual output, *y* of the controlled plant. Note that in drawing an analogy with the eye-blink conditioning paradigm, we use the simplifying convention of considering the noxious stimulus (the air-puff) as a reference, *r*, that indicates that the eyelids should close; the closure of the eyelid as the output of the plant, *y*; and the sensory response to the noxious stimulus as an error, *e*, that encodes the difference between the desired, *r*, and the actual eyelid closures, *y*. Given this, we advance a new learning rule, *f*, that achieves optimal performance in the context of eye-blink conditioning and other cerebellar learning paradigms.

### 2.2 Cerebellar Control Architecture

We embed the adaptive filter cerebellar module in a layered control architecture based on the interaction between brain stem motor nuclei driving motor reflexes and the cerebellum, such as the one established between the cerebellar microcircuit responsible for conditioned responses and the brain stem reflex circuitry that produces unconditioned eye-blinks [Christian and Thompson, 2003] (Fig. 2 *left*). In our model, we assume that cerebellar output, *o*, can be interpreted as a signal coded in the sensory domain that feeds the lower reflex controller (Fig. 2 *right*). Abstracting from the anatomical layout, we formulate a layered control architecture, the CFPC, in which an adaptive feed-forward layer supplements a negative feedback controller, steering it with feed-forward signals.

**Figure 2:**
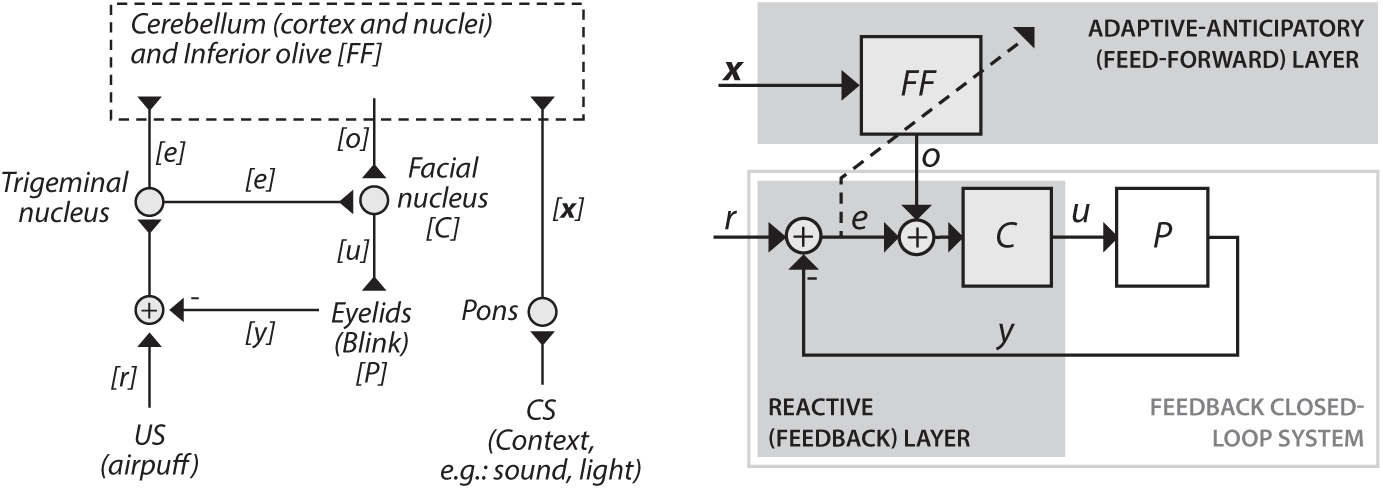
Neuroanatomy of eye-blink conditioning and the CFPC architecture. *Left*: Mapping of signals to anatomical structures in eye-blink conditioning [De Zeeuw and Yeo, 2005, Medina et al., 2002]; regular arrows indicate inputs and outputs that interact in the world, arrows with inverted heads indicated neural pathways. *Right*: CFPC architecture. Abbreviations: *P*, plant; *C*, feedback controller; *FF*, feed-forward module; *r*, reference signal; *y*, plant’s output; *e*, output error; **x**, basis signals; *o*, feed-forward signal; and *u*, motor command.

Our architecture uses a single-input single-output negative feedback controller. The controller receives as input the output error *e*, that is the difference between a reference signal *r* and the output of the plant *y*. For the derivation of the learning algorithm, we only assume that both plant and controller are linear and time-invariant (LTI) systems. Importantly, the feedback controller and the plant form a reactive closed-loop system, that mathematically can be seen as a system that maps the reference, *r*, into the plant’s output, *y*. A feed-forward layer that contains the above-mentioned cerebellar model provides the negative feedback controller with an additional input signal, *o*. We refer to *o* as a *counter-factual* error signal, since although it *mechanistically* drives the negative feedback controller analogously to an error signal it is not an *actual* error. The counterfactual error is generated by the feed-forward module that receives an output error, *e*, as its teaching signal. Notably, from the point of view of the reactive layer closed-loop system, *o* can also be interpreted as a signal that offsets *r*. In other words, even if *r* remains the reference that sets the target of behavior, *r* + *o* functions as the *effective* reference that drives the closed-loop system.

## 3 Results

### 3.1 Derivation of the gradient descent update rule for the cerebellar control architecture

We have designed a cerebellar adaptive-anticipatory control architecture (Fig. 2). The architecture is built upon a reflex, modeled as an output error feedback controller that controls a given plant. The architecture contains a feed-forward module whose output signal, added to the output error, enters the feedback controller (i.e, triggering the reflex). The feed-forward module acts as a linear filter combining a set of *G* bases. The linear combination of the bases is updated based on the same error information that drives the feedback controller. The task of the overall architecture is to follow a finite reference signal **r** ϵ ℝ^*T*^ that is repeated trial-by-trial. To analyze this system, we will use the discrete time formalism and assume that all components are linear time-invariant (LTI). Given this, both reactive controller and plant can be lumped together into a closed-loop dynamical system, that can be described with the dynamics **A**, input **B**, measurement **C** and feed-through **D** matrices. In general, these matrices describe how the state of a dynamical system autonomously evolves with time, **A**; how inputs affect system states, **B**; how states are mapped into outputs, **C**; and how inputs instantaneously affect the system’s output **D**. As we consider a reference of a finite length *N*, we can construct the *N*-by-*N* transfer matrix 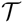 as follows:

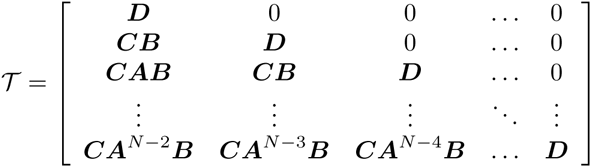

With this transfer matrix we can map any given reference **r** into an output **y** using **y** = 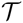**Tr**, obtaining the complete output trajectory of an entirely feedback-driven trial. Note that the first column of 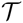 contains the impulse response curve of the closed-loop system, while the rest of the columns are obtained shifting that impulse response by *n* – 1 time steps, with *n* indexing the column. Therefore, we can build the transfer matrix 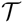 either in a model-based manner, deriving the state-space characterization of the closed-loop system, or in measurement-based manner, measuring the impulse response curve. Additionally, note that (**I** – 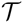)**r** yields the error of the feedback control in following the reference *r*, which we denote with **e**_0_.

Let **o** ϵ ℝ^*N*^ be the entire feed-forward signal for a given trial. Given commutativity, we can consider that from the point of view of the closed-loop system, it is added directly to the reference, *r*, (Fig. 2 *right*). In that case, we can use **y** = 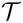 (**r** + **o**) to obtain the output of the closed-loop system when it is driven by both the reference and the feed-forward signal, **o**. Note that we can resolve for the optimal feed-forward signal (**o**^∗^) that will cancel the error with 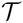^*†*^**e**_0_, using the Moore-Penrose pseudo-inverse (†). However, rather than being able to generate any arbitrary output signal, the feed-forward module only outputs linear combinations of a set of bases. Let **X** ϵ ℝ^*N* × *G*^ be a matrix with the content of the *G* bases during *N* time steps. The feed-forward signal becomes **o** = **Xw**, where **w** ϵ ℝ^*G*^ contains the mixing weights. Hence, the output of the architecture given a particular **w** becomes **y** = 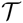(**r** + **Xw**).

We implement learning as the process of adjusting the weights **w** of the feed-forward module. It occurs in a trial-by-trial manner, such that at each trial the same reference signal (**r**) and bases (**X**) will be repeated. Through learning we want to converge to the optimal weight vector **w**^∗^ defined as

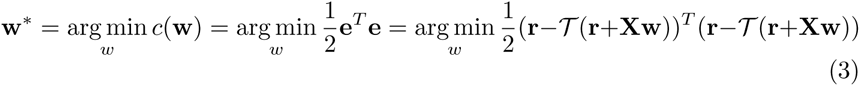

where *c* indicates the objective function to minimize, namely the *L*_2_ norm or sum of squared errors. With the substitution 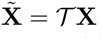 and using **e**_0_ = (**I** – 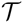)**r**, the minimization problem can be cast as a *canonical* linear least-squares problem:

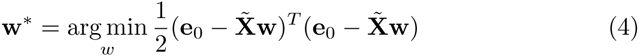

One the one hand, this allows to directly find the least squares solution for **w**^∗^, that is, 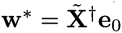. On the other hand, and more interestingly, with **w**[*k*] being the weights at trial *k* and having 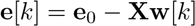, we can obtain the gradient of the error function at trial *k* with relation to ***w*** as follows:

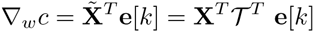

Thus, setting *η* as a properly scaled learning rate (the only global parameter Θ of the rule), we can derive the following gradient descent strategy for the update of the weights between trials:

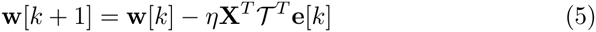

This solves for the learning rule *f* in eq. 2. Note that *f* is consistent with both the cerebellar anatomy (Fig. 2*left*) and the control architecture (Fig. 2*right*) in that, in addition to the basis inputs, the feed-forward module/cerebellum only requires the error signal to update the weights/efficacies.

### 3.2 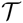^*T*^ plays the role of a synaptic eligibility trace

The standard least mean squares (LMS) rule (also known as Widrow-Hoff or decorrelation learning rule) can be represented in its batch version as **w**[*k* + 1] = **w**[*k*] – *η***X**^*T*^ **e**[*k*]. Hence, the only difference between the batch LMS rule and the one we have derived is the insertion of the matrix factor 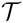^*T*^. Now we will show how this factor acts as a filter that computes an eligibility trace at each synapse/weight. Note that the update of a single weight, according Eq. 5 becomes

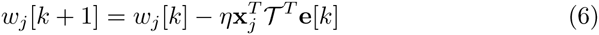

where **x**_*j*_ contains the sequence of values of the cortical basis *j* during the entire trial. This can be rewritten as

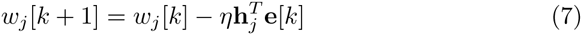

with **h**_*j*_ ≡ 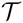**x**_*j*_. The above inner product can be expressed as a sum of scalar products

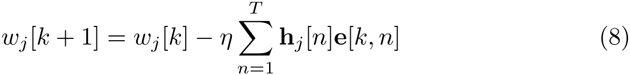

where *n* indexes the within trial time-steps. Note that **e**[*k*] in eq. 7 refers to the whole error signal at trial *k* whereas **e**[*k, n*] in eq. 8 refers to the error value in the *n*-th time-step of the trial *k*. It is now clear that each component *h*_*j*_ weighs how much an error arriving at time *n* should modify the weight *w*_*j*_, which is precisely the role of an eligibility trace. Note that since 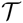 contains in its columns/rows shifted repetitions of the impulse response curve, the eligibility trace codes at any time *n*, the convolution of the sequence of previous inputs with the impulse-response curve of the reactive layer closed-loop. Indeed, in each synapse, the eligibility trace is generated by a forward model of the closed-loop system that is exclusively driven by the basis signal.

Consequently, our main result is that by deriving a gradient descent algorithm for the cerebellar control architecture introduced above we have obtained an exact definition of the eligibility trace. That definition guarantees that the set of weights/synaptic efficacies are updated in a manner locally optimal in the weights’ space.

### On-line gradient descent algorithm

The trial-by-trial formulation above allowed for a straightforward derivation of the (batch) gradient descent algorithm. As it lumped together all computations occurring in a same trial, it accounted for time within the trial implicitly rather than explicitly: one-dimensional time-signals were mapped onto points in a high dimensional space. However, after having established the gradient descent algorithm, we can implement the same rule in an on-line manner, dropping the trial-by-trial assumption and performing all computations locally in time. Each weight/synapse must have a process associated to it that outputs the eligibility trace. That process passes the incoming (unweighted) basis signal through a (forward) model of the closed-loop as follows:

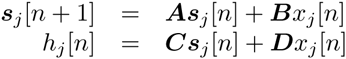

where matrices ***A***, ***B***, ***C*** and ***D*** refer to the closed-loop system (they are the same matrices that we used to define the transfer matrix 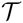), and ***s***_*j*_[*n*] is the state vector of the forward model of the synapse *j* at time-step *n*. In practice, each “synaptic” forward model computes what would have been the effect of having driven the closed-loop system with each basis signal alone. Given the superposition principle, the outcome of that computation can also be interpreted as saying that *h*_*j*_[*n*] indicates what would have been the displacement over the current output of the plant, *y*[*n*], achieved using the unweighted basis signal *x*_*j*_. The process of weight update is completed as follows:

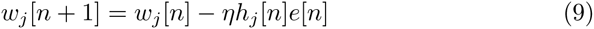

At each time step *n*, the error signal *e*[*n*] is multiplied by the current value of the eligibility trace *h*_*j*_[*n*], scaled by the learning rate *η*, and subtracted to the current weight*w*_*j*_[*n*]. Therefore whereas the contribution of each basis to the current output of the adaptive filter depends only on its current value and weight, the change in weight depends on the current and past values passed through a forward model of the closed-loop dynamics.

### 3.4 Simulation of a visually-guided smooth pursuit task

We demonstrate the CFPC approach in an example of a visual smooth pursuit task in which the eyes have to track a target moving on a screen. Even though the simulation does not capture all the complexity of a smooth pursuit task, it illustrates our anticipatory control strategy. We model the plant (eye and ocular muscles) with a two-dimensional linear filter that maps motor commands into angular positions. Our model is an extension of the model in [Porrill and Dean, 2007], even though in that work the plant was considered in the context of the vestibulo-ocular reflex. In particular, we use a chain of two leaky integrators: a slow integrator with a relaxation constant of 100 ms drives the eyes back to the rest position; the second integrator, with a fast time constant of 3 ms ensures that the change in position does not occur instantaneously. To this basic plant, we add a reactive control layer modeled as a proportional-integral (PI) error-feedback controller, with proportional gain *k*_*p*_ and integral gain *k*_*i*_. The control loop includes a 50 ms delay in the error feedback, to account for both the actuation and the sensing latency. We choose gains such that reactive tracking lags the target by approximately 100 ms. This gives *k*_*p*_ = 20 and *k*_*i*_ = 100. To complete the anticipatory and adaptive control architecture, the closed-loop system is supplemented by the feed-forward module. The architecture implementing the forward model-based gradient descent algorithm is applied to a task structured in trials of 2.5 sec duration. Within each trial, a target remains still at the center of the visual scene for a duration 0.5 sec, next it moves rightwards for 0.5 sec with constant velocity, remains still for 0.5 sec and repeats the sequence of movements in reverse, returning to the center. The cerebellar component receives 20 Gaussian basis signals (**X**) whose receptive fields are defined in the temporal domain, with a width (standard-deviation) of 50 ms and spaced by 100 ms. The whole system is simulated using a 1 ms time-step. To construct the matrix 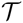 we computed closed-loop system impulse response.

At the first trial, before any learning, the output of the plant lags the reference signal by approximately 100 ms converging to the position only when the target remains still for about 300 ms (Fig. 3 *left*). As a result of learning, the plant’s behavior sifts from a reactive to an anticipatory mode, being able to track the reference without any delay. Indeed, the error that is sizable during the target displacement before learning, almost completely disappears by the 50^*th*^ trial (Fig. 3 *right*). That cancellation results from learning the weights that generate a feed-forward predictive signal that leads the changes in the reference signal (onsets and offsets of target movements) by approximately 100 ms (Fig. 3 *right*). Indeed, convergence of the algorithm is remarkably fast and by trial 7 it has almost converged to the optimal solution (Fig. 4).

**Figure 3:**
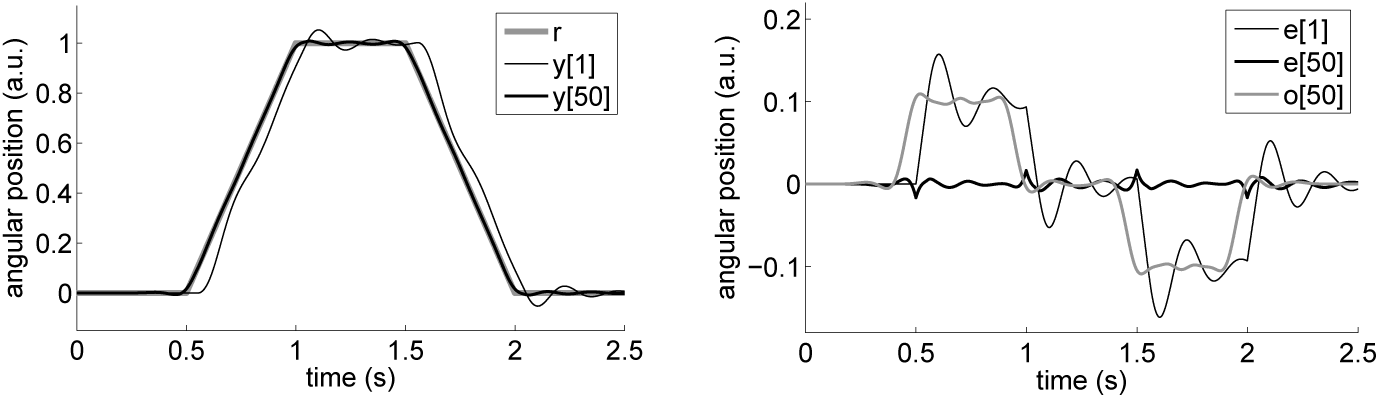
Left: Reference (**r**) and output of the system before (**y**[1]) and after learning (**y**[50]). Right: Error before **e**[1] and after learning **e**[50] and output acquired by cerebellar/feed-forward component (**o**[50])

**Figure 4:**
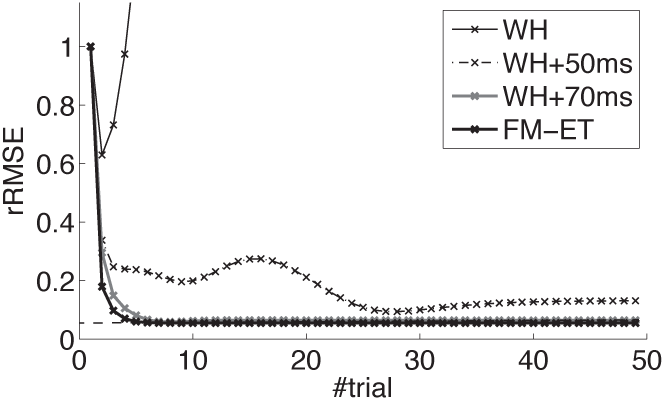
Representative learning curves of the forward model-based eligibility trace gradient descent (FM-ET), the simple Widrow-Hoff (WH) and the Widrow-Hoff algorithm with a delta-eligibility trace matched to error feedback delay (WH+50 ms) or with an eligibility trace exceeding that delay by 20 ms (WH+70 ms). Error is quantified as the relative root mean-squared error (rRMSE), scaled proportionally to the error in the first trial. Error of the optimal solution, obtained with 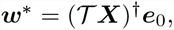 is indicated with a dashed line

To assess how much our forward-model-based eligibility trace contributes to performance, we test two alternative algorithms. In both cases we employ the same control architecture, changing the plasticity rule such that we either use no eligibility trace, thus implementing the basic Widrow-Hoff learning rule, or use the Widrow-Hoff rule extended with a delta-function eligibility trace that matches the latency of the error feedback (50 ms). Performance with the basic WH model worsens rapidly whereas performance with the WH learning rule using a “pure delay” eligibility trace improves but not as fast as with the forward-model-based eligibility trace (Fig. 4). Indeed, the best strategy for implementing delayed delta eligibility traces is to set a delay exceeding the transport delay by around 20 ms, thus matching the peak of the impulse response. In that case, the system performs almost as good as with the forward-model eligibility trace (70 ms). This last result implies that, even though the literature usually emphasizes the role of transport delays, eligibility traces also account for response lags due to intrinsic dynamics of the plant.

To summarize our results, we have shown with a basic simulation of a visual smooth pursuit task that generating the eligibility trace by means of a forward model ensures convergence to the optimal solution accelerates learning by guaranteeing that it follows a gradient descent.

## Discussion

The first seminal works of cerebellar computational models emphasized its role as an associative memory [Marr, 1969, Albus, 1971]. Later, [Fujita, 1982] and [Dean et al., 2010] investigated the cerebellum as a device processing correlated time signals. In this latter framework, the use of the computational concept of an eligibility trace emerged as a heuristic construct that allowed to compensate for the transmission delays in the circuit, which introduced lags in the cross-correlation between signals. Concretely, that was referred as the problem of *delayed error feedback* due to which by the time an error signal reaches a cell, the synapses accountable for that error are not the ones currently active, but those that were (or could have been) active at the time when the motor signals that caused the actual error were generated. This view however neglected the fact that beyond transport delays, the response dynamics of physical plants also influence how the past pre-synaptic signals could have related to the current output of the plant. Indeed, for a linear plant, the impulse-response function of the plant provides the complete description of how inputs will drive the system, and as such, integrates transmission delays as well as the dynamics of the plant.

In cerebellar control models discussed in [Kettner et al., 1997] and [Porrill and Dean, 2007] the temporal misalignment between error feedback and synaptic activity was dealt with as a misalignment between temporal signals. This led to the intuition that a delta-function eligibility trace was an effective mechanism for functionally realigning those signals. However, as exact delayed-response delta-function filters in Purkinje cell synapses were not considered biologically realistic, broader alpha-function eligibility traces were proposed (also in [Kettner et al., 1997]) as a biologically-plausible mechanism implementing an approximation to a delay, whereby one could keep a temporal memory of pre-synpatic inputs. Our frame-work refines that conclusion: both broad or narrow eligibility traces lead to near-optimal performance depending upon the system’s dynamics. Notably, in our model we observe that an exact delta-function eligibility trace is optimal only for a trivial plant with no dynamics.

Here, with the CFPC, we have modeled the cerebellar system at a very high level of abstraction: we have not included bio-physical constraints underlying neural computations, obviated known anatomical connections such as the cerebellar nucleo-olivary inhibition [Bengtsson and Hesslow, 2006, Herreros and Verschure, 2013] and made simplifications such as collapsing cerebellar cortex and nuclei into the same computational unit. On the one hand, such a choice of high-level abstraction may indeed be beneficial for deriving general-purpose machine learning or adaptive control algorithms. On the other hand, it is remarkable that in spite of this abstraction our framework makes fine-grained predictions at the micro-level of biological processes. In conclusion, our systems level CFPC model of cerebellar computation provides a normative interpretation of plasticity in Purkinje cell synapses: in order to generate optimal eligibility traces, synapses must include a forward model of the controlled subsystem.

